# Establishment of Rabbit Formative-like Pluripotent Cells by Bidirectional Modulation of WNT Signaling Pathway

**DOI:** 10.64898/2026.10.13.757547

**Authors:** Yuanxi Yang, Manya Yu, Tong Qiu, Le Zheng, Tao Cui, Zhengpeng Zhuang, Lei Li, Longquan Quan, Junjie Mao, Liangxue Lai

## Abstract

The generation of embryonic stem cells (ESCs) with full developmental capacity and robust chimera-forming potential remains a major bottleneck for most non-rodent mammalian species, including rabbits, an important animal for both biomedicine and agriculture. Formative pluripotency is a unique intermediate state bridging the naive and primed pluripotent states, which is defined by distinct molecular signatures, intrinsic competence for primordial germ cell (PGC) specification, and unique developmental adaptability to early embryonic microenvironments. Here, we developed a novel culture medium termed KFiC, which could bidirectionally and coordinately modulate the WNT/β-catenin signaling pathway via combined treatment of the WNT activator CHIR99021 and the inhibitor IWR-1. The KFiC culture system enables efficient derivation and stable long-term self-renewal of rabbit embryonic stem cells (rbESCs) with consistent formative-like pluripotent characteristics. KFiC-cultured rbESCs (KFiC-rbESCs) form compact, undifferentiated colonies, maintain intact genomic stability, and recapitulate the transcriptional landscape of pre-implantation epiblast cells, accompanied by sustained high expression of the key formative pluripotency marker OTX2. Functionally, KFiC-rbESCs possess efficient PGC differentiation competence and extensively contribute to embryonic development after microinjection into rabbit morulae, displaying their superior developmental plasticity. Notably, the KFiC system can stabilize an intermediate formative-like pluripotent state in mouse, human, and pig pluripotent stem cells, and KFiC-adapted mouse ESCs retain potent chimera-forming ability. Collectively, this study establishes a reliable culture platform for generating and maintaining formative-like rbESCs. This cross-species-compatible system provides a promising foundation for further optimizing naïve-level pluripotency in rbESCs and facilitates the generation of genetically engineered rabbit disease models.

## Introduction

The establishment of authentic embryonic stem cells (ESCs) capable of long-term self-renewal while faithfully retaining embryonic developmental potential remains a major challenge for many mammalian species. The rabbit (Oryctolagus cuniculus) is an important agricultural and experimental model that shares several physiological and developmental features with humans and has been widely used in biomedical research, including disease modeling, regenerative medicine, and developmental studies^1–3^. However, despite longstanding efforts to establish rabbit ESCs (rbESCs) with germline competency comparable to that of mouse ESCs, the derivation and long-term maintenance of such cells have posed substantial challenges. Yet recent advances have begun to expand the developmental potential of rabbit pluripotent stem cells. Several culture systems, including KF^4^, rESM^5–7^ and bXC^8^, have been explored for the derivation and expansion of rbESCs, but the resulting cells have generally displayed restricted pluripotency and developmental potential. More recently, Wang et al. developed 6iL/E4, a culture system that supports the maintenance of rbESCs and enable chimeric contribution following their injection into rabbit morulae, but germline transmission was not established^9^. In parallel, Pham et al. claimed that overexpression of KLF2, ERAS, and PRMT6 (KEP), together with an optimized culture medium, conferred germline competency on rabbit induced pluripotent stem cells (rbiPSCs). However, germline transmission of rbiPSC-derived germ cells through the resulting F0 animals was not described^10^.

Of note, WNT signaling is unidirectionally modulated to varying degrees in many of above culture media, highlighting the potential importance of this pathway in determining rabbit pluripotency. The WNT signaling pathway plays extensive roles in many biological processes, including cell fate specification, tissue development, and stem cell self-renewal, and its regulatory function in pluripotency is highly context- and species-dependent, with distinct pluripotent states requiring different levels of WNT activity^11^. In mouse naïve ESCs, activation of canonical WNT signaling through GSK3 inhibition by CHIR99021 promotes pluripotency and self-renewal^12^, whereas in human naïve pluripotent stem cells, WNT signaling is generally suppressed through inhibition of the pathway by XAV939^13^. Interestingly, bidirectional modulation of WNT signaling through combined treatment with the WNT activator CHIR99021 and the WNT inhibitor XAV939 has been shown to support the maintenance of primed pluripotency in both mouse and human ESCs^14^.

In addition to naïve and primed states, accumulating evidence has established formative pluripotency as a distinct and transient phase that emerges between naïve pluripotency and the primed epiblast during peri-implantation development^15–17^. Formative pluripotent cells possess a unique molecular program characterized by activation of OTX2^18^, a key transcriptional regulator of this stage, together with progressive establishment of de novo DNA methylation mediated by DNMT3A and DNMT3B^19^ and metabolic remodeling to glycolysis^20^. Functionally, formative pluripotency is associated with competence for primordial germ cell (PGC) specification in response to inductive signals directly, providing a defined route for generating PGC-like cells (PGCLCs) in vitro^21,22^. Importantly, formative ESCs also retain substantial developmental potential in vivo and can contribute to the germline when introduced into early-stage embryos^17^. Thus, culture media that capture and stabilize formative pluripotency may provide an effective strategy for expanding the developmental potential of ESCs in species in which a conventional naïve state has proven difficult to establish and maintain.

Despite considerable efforts, previously established rbESCs have generally exhibited a primed pluripotent state resembling mouse epiblast stem cells (EpiSCs), accompanied by limited developmental potential^4,23–25^. Here, we developed KFiC, a WNT-modulated culture medium based on the KF system that enables the efficient establishment and long-term maintenance of rabbit ESCs with formative-like features through bidirectionally coordinated regulation of WNT signaling using the activator CHIR99021 and the inhibitor IWR-1. In our results, rbESCs maintained in KFiC (KFiC-rbESCs) display stable self-renewal capacity, preserved genomic integrity, and sustained robust expression of formative-associated genes, while their global transcriptional profiles closely resembled those of pre-implantation epiblast populations. Functionally, KFiC-rbESCs retained competence for PGC induction and contributed to embryonic chimerism following introduction into rabbit morulae. Moreover, KFiC supports an intermediate pluripotent state in ESCs from multiple mammalian species, including mouse, pig, and human. Together, our findings establish a WNT-regulated culture system for the efficient derivation and long-term maintenance of formative-like rabbit ESCs and provide evidence that coordinated WNT modulation can promote a reversible intermediate pluripotent state across many mammalian species.

## Results

### Bidirectional modulation of the WNT pathway enables efficient derivation and long-term maintenance of rbESCs

Since embryonic stem cells (ESCs) have been established from blastocysts in numerous mammalian species, with multiple culture systems developed within individual species to support distinct pluripotent states, we first tested these previously reported PSC culture conditions for their ability to support rabbit ESCs (Fig. 1A). Human naïve PSC media, such as 4CL^26^ and PXGL^27^, failed to support the formation of outgrowths from rabbit blastocysts under our experimental conditions.

**Figure 1.**
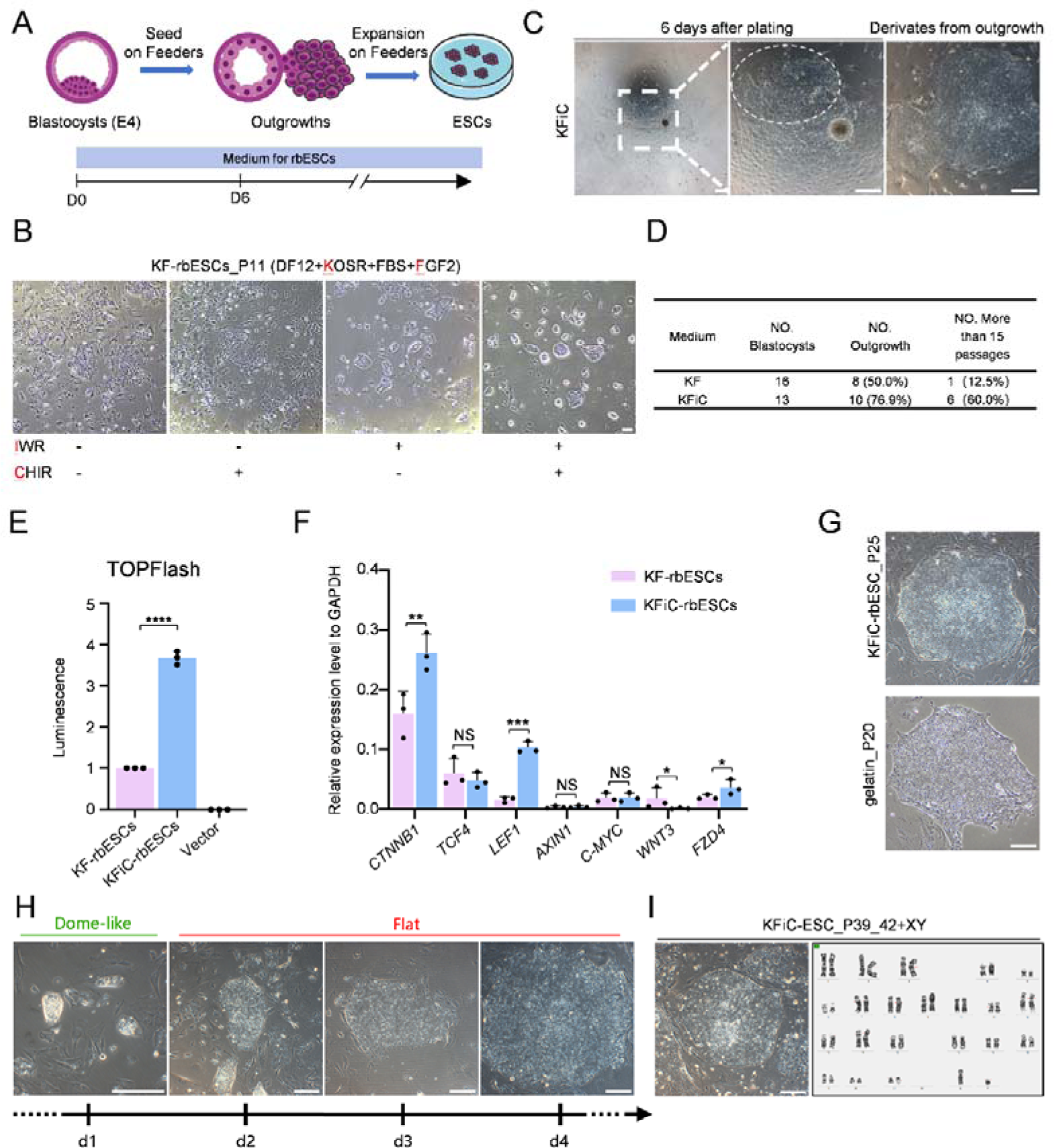
Bidirectional Modulation of the WNT Signaling Enables Efficient Derivation and Long-Term Maintenance of Rabbit ESCs. (**A**) Schematic diagram of rbESC isolation. E4.0 blastocysts with intact zona pellucida were plated onto feeder cells in basal medium supplemented with small molecules. Adherent blastocysts were cultured without agitation until clear outgrowths emerged. Compact outgrowths were then mechanically isolated, dissociated, and replated onto feeder cells for further proliferation. (**B**) Effects of IWR-1 and CHIR99021 on P8 KF-rbESCs colony formation over three passages. IWR, IWR-1; CHIR, CHIR99021. Scale bar, 200 μm. (**C**) Representative morphology of rabbit outgrowths and their derivatives cultured in KFiC medium during rbESC derivation. Scale bar, 200 μm. (**D**) Summary of rbESC derivation efficiency in KF and KFiC media. (**E**) Evaluation of WNT signaling activity using the TOPFlash reporter assay. Data represent the mean ± SD from three independent experiments (n = 3). *P < 0.05; **P < 0.01; ***P < 0.001; ****P < 0.0001. (**F**) RT-qPCR analysis of WNT signaling-related genes in rabbit ESCs cultured in KF and KFiC. Data represent the mean ± SD from three independent experiments (n = 3). *P < 0.05; **P < 0.01; ***P < 0.001. (**G**) Morphology of rbESCs during long-term culture in KFiC medium. Top, cells grown on feeder cells; Bottom, cells grown under feeder-free conditions on gelatin-coated dishes. Scale bar, 200 μm. (**H**) Colony morphology of KFiC-rbESCs over successive passages following single-cell dissociation with trypsin. Scale bar, 200 μm. (**I**) Characterization of rbESCs during long-term culture. Left, representative colony morphology; right, karyotype analysis. Scale bar, 200 μm.

In contrast, 2i/LIF^12^ and FTW^28^, developed for mouse PSCs, facilitated early expansion of rabbit blastocyst-derived outgrowths but were insufficient to sustain their long-term self-renewal (Fig. S1A, S1B). Notably, FTW induced the emergence of dome-shaped outgrowths, a morphological feature of mouse naïve ESCs, and this phenotype was further enhanced by supplementation with 5% KOSR. However, the enhanced phenotype was transient and persisted only during the first three passages, after which the cultured cells underwent rapid differentiation and death (Fig. S1C). Consistent with these findings, FACE, a culture system designed for ruminant animals that also contains 3 μM CHIR99021^29^, similarly failed to support the long-term self-renewal of rbESCs (Fig. S1D). We subsequently evaluated T2iL, a modified 2i/LIF medium developed for rat ESCs by reducing the GSK3 inhibitor concentration from 3 μM to 1 μM and reported to provide a more robust platform for ESC expansion^30,31^. In our data, although T2iL supported the derivation of rabbit embryonic outgrowths, it failed to sustain rbESCs during long-term culture (Fig. S1E).

Multiple media including 2i/LIF, FTW, FACE, and T2iL promoted early expansion of rabbit blastocyst-derived outgrowths and all involved activation of WNT pathway signaling. we therefore hypothesized that fine-tuned WNT/β-catenin signaling is critical for rbESC self-renewal. To test this hypothesis, we modified KF medium (AKSF: Accutase, KOSR, FCS, and FGF), which supports rbESC derivation, single-cell dissociation, and rapid proliferation but fails to support their contribution to the inner cell mass (ICM) of pre-implantation embryos^4^. We found that, although KF medium initially enabled colony formation and expansion of rbESCs (KF-rbESCs), most colonies progressively lost their compact morphology upon extended passaging (Fig. S1F). We therefore sought to supplement KF with CHIR99021, a GSK3 inhibitor that activates canonical WNT signaling by preventing phosphorylation-mediated degradation of β-catenin^32^, together with IWR-1, a tankyrase inhibitor that promotes β-catenin degradation and limits its nuclear accumulation and thereby attenuates excessive WNT activation, to achieve coordinated modulation of WNT/β-catenin signaling in this system^33^.

We first supplemented KF medium with titrated CHIR99021 or IWR-1 individually, but neither treatment alone prevented the progressive loss of compact colony morphology (Fig. 1B). In contrast, KF medium supplemented with titrated CHIR99021 and IWR-1, termed KFiC, markedly improved colony morphology, resulting in more compact colonies with smoother and better-defined boundaries within only three passages (Fig. 1B). We then evaluated the capacity of KFiC to support the de novo establishment of rbESC lines from blastocysts (Fig. 1A). KFiC supported outgrowth formation from 76.9% of blastocysts (10/13), compared with 50.0% (8/16) under KF conditions. Of the lines subsequently established, 6 of 10 (60%) established in KFiC were maintained beyond passage 15 while retaining compact colony morphology, whereas only 1 of 8 (12.5%) under KF conditions reached this criterion (Fig. 1 D). Together, these findings demonstrate that KFiC enhances both the initial outgrowth formation and subsequent long-term maintenance of blastocyst-derived rbESC lines.

Consistent with our hypothesis, TOPFlash reporter assays revealed that canonical WNT activity was increased by more than threefold in KFiC compared with KF, accompanied by elevated expression of key WNT pathway components, including *CTNNB1* and *LEF1* (Fig. 1E, F). Moreover, rbESCs cultured in KFiC (KFiC-rbESCs) proliferated robustly during long-term culture under both feeder-supported and feeder-free conditions (Fig. 1G), while undergoing dynamic changes in colony morphology following single-cell dissociation. Specifically, KFiC-rbESCs initially formed compact, dome-shaped colonies within 24 h and subsequently transitioned into flattened EpiSC-like colonies over the following 72 h (Fig. 1H). In addition, KFiC-rbESCs retained a stable diploid karyotype after more than 39 consecutive passages, with the expected complement of 44 chromosomes, comprising 42 autosomes and XY sex chromosomes (Fig. 1I). Collectively, these findings demonstrate that KFiC-mediated WNT modulation enables efficient rbESC derivation and long-term maintenance with compacted colony morphology, robust proliferation, and genomic stability.

### KFiC sustains a pre-implantation epiblast-like state in rabbit ESCs

To evaluate the pluripotency properties, we first investigated the expression of core pluripotency-associated factors. Immunostaining and western blot analyses showed higher and more stable expression of OCT4 and SOX2 in KFiC-rbESCs than in KF-rbESCs; more problematically, their expression in KF-rbESCs was reduced or undetectable across independent experiments (Fig. 2A, 2B). Additionally, KFiC-rbESCs displayed strong alkaline phosphatase (AP) activity and readily formed embryoid bodies in suspension culture conditions (Fig. 2C, D). Furthermore, following subcutaneous transplantation into immunodeficient mice, KFiC-rbESCs gave rise to well-developed teratomas (Fig. S2A) containing derivatives of all three germ layers (Fig. 2E).

**Figure 2.**
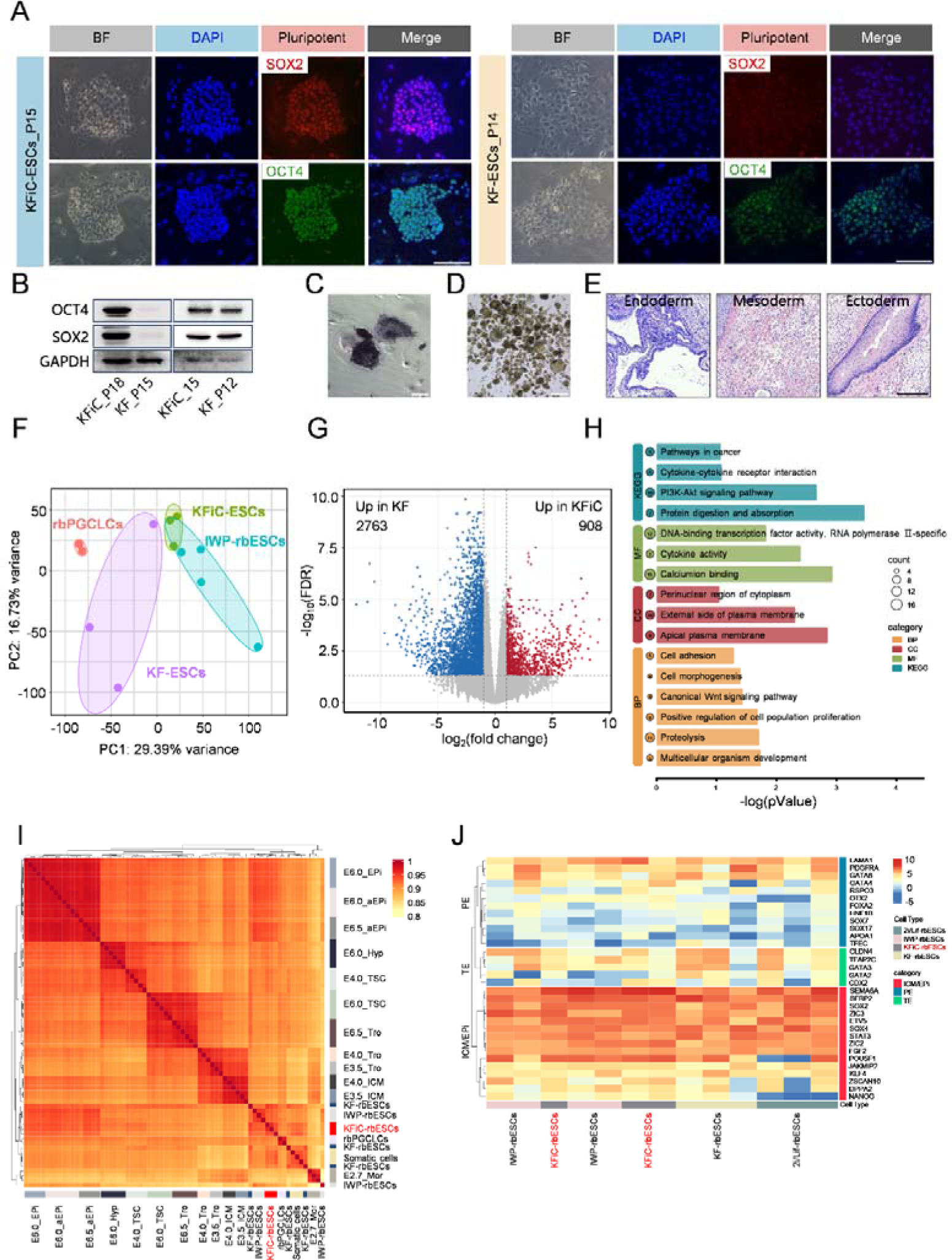
Pluripotent State of Rabbit ESCs Cultured in KFiC. (**A**) Immunofluorescence detection of pluripotency markers SOX2 and OCT4 in rbESCs. Left, KFiC medium; right, KF medium. Nuclei were counterstained with DAPI. Scale bar, 200 μm. (**B**) Western blot analysis of pluripotency markers SOX2 and OCT4 in rbESCs cultured in KFiC or KF medium. GAPDH was used as a loading control. (**C**) Alkaline phosphatase (AP) staining of P15 KFiC-rbESCs. Scale bar, 200 μm. (**D**) Embryoid body formation after 7 days of suspension culture in DMEM containing 10% FBS. Scale bar, 200 μm. (**E**) Representative H&E-stained sections of teratoma displaying derivatives of the ectoderm, mesoderm, and endoderm. Scale bar, 200 μm. (**F**) Principal component analysis (PCA) of the transcriptomes of KFiC-rbESCs, KF-rbESCs, IWP-rbESCs, and rbPGCLCs. RNA-seq data for all expressed genes were used for the analysis. (**G**) Differentially expressed genes between KFiC and KF (FC > 2). (**H**) Gene Ontology (GO) enrichment analysis of the top 200 differentially expressed genes (DEGs) between KFiC-rbESCs and KF-rbESCs. Significantly enriched terms were defined using a Benjamini–Hochberg adjusted p-value < 0.05. (**I**) Hierarchical clustering dendrogram showing transcriptomic distances among samples. (**J**) Clustered heatmap of lineage-enriched genes across rbESC lines. Genes enriched in ICM/epiblast, PE, and TE are shown for KFiC-rbESCs, KF-rbESCs, 2i/LIF-rbESCs, and IWP-rbESCs.

Next, we performed RNA-seq analysis of KFiC-rbESCs, KF-rbESCs, 2i/LIF-rbESCs (rbESCs cultured in 2i/LIF), and together with previously reported rabbit pluripotent populations (IWP-rbESCs, rbESCs cultured in IWP; rbPGCLCs, rabbit primordial germ cell-like cells)^34^ to characterize their molecular identities. Principal component analysis revealed distinct transcriptional profiles among these samples, with KFiC-rbESCs forming a relatively compact cluster, indicative of greater transcriptional consistency among biological replicates (Fig. 2F). Differential expression analysis (Log2 fold change, >2; adjusted p < 0.05) identified 908 genes upregulated in iKFC-rbESCs cells relative to KF-rbESCs (Fig. 2G). Gene Ontology (GO) enrichment analysis showed that DEGs between KFiC-rbESCs and KF-rbESCs were enriched in terms related to cytokine activity and multicellular organismal development. In contrast, DEGs between KFiC-rbESCs and IWP-rbESCs were enriched in terms associated with integrin binding and regulation of cell migration (Fig. 2H; Fig. S2C).

Having identified the distinct transcriptional profile of KFiC-rbESCs, we next sought to position them along the rabbit developmental continuum by comparing their transcriptomic profiles with a publicly available rabbit embryonic reference dataset spanning successive developmental stages^35^. Hierarchical clustering placed KFiC-rbESCs closest to rbPGCLCs (Fig. 2I, Fig. S2C), which have been associated with a pre-implantation epiblast-like state in rabbits^34^. Analysis of lineage-specific genes further showed that KFiC-rbESCs expressed higher levels of ICM/epiblast-associated genes, such as *SEMA6A*, *ZIC3*, and *ZSCAN10* (Fig. 2J). Whereas KF-rbESCs exhibited elevated expression of trophoblast-associated markers, including *TFAP2C*, *GATA2*, and *GATA3*, and 2i/LIF-rbESCs displayed diminished expression of key pluripotency regulators, including *POU5F1* and *NANOG*, patterns consistent with their limited long-term self-renewal and their progressive differentiation during extended culture (Fig. 2J; Fig. S1B).

Collectively, these findings position KFiC-rbESCs within a pre-implantation epiblast-like transcriptional state, underscoring the importance of bidirectional WNT modulation in maintaining a stable pluripotent state in rabbit ESCs.

### KFiC stabilizes a formative-like pluripotent state in rbESCs

Based on our transcriptomic analysis, we found that KFiC-rbESCs shared transcriptional similarities with pre-implantation epiblast-like cells and E6.5 anterior epiblasts (Fig. 2I, Fig. S2D), corresponding to the peri-implantation developmental window in rabbits^4,36^. These findings suggested that KFiC-rbESCs may stay in a formative-like pluripotent state. We therefore examined the expression profiles of genes corresponding to naïve, formative, primed, and primitive streak stages.

In KFiC-rbESCs, we observed that naïve-associated markers, including *OOEP*, *DPPA5*, and *KHDC3L*, were moderately expressed (Fig. 3A and Table S1), whereas classical primed-state genes, such as *LEFTY1/2* and *NODAL*, showed limited expression. To define the formative identity, we evaluated the expression of *OTX2*, a key regulator of formative pluripotency^17^, and found that it was highly expressed in KFiC-rbESCs (Fig. 3A, B). Consistently, immunostaining and western blot analyses confirmed elevated OTX2 protein levels in KFiC-rbESCs compared with KF-rbESCs (Fig. 3C, D). Furthermore, staining of ZO-1, a tight junction-related protein that delineates cell boundaries^37^, revealed a higher nuclear-to-cytoplasmic ratio in KFiC-rbESCs than in KF-rbESCs (Fig. 3E, F). This morphological feature, together with the expression of OTX2, supports the formative-like identity of KFiC-rbESCs.

**Figure 3.**
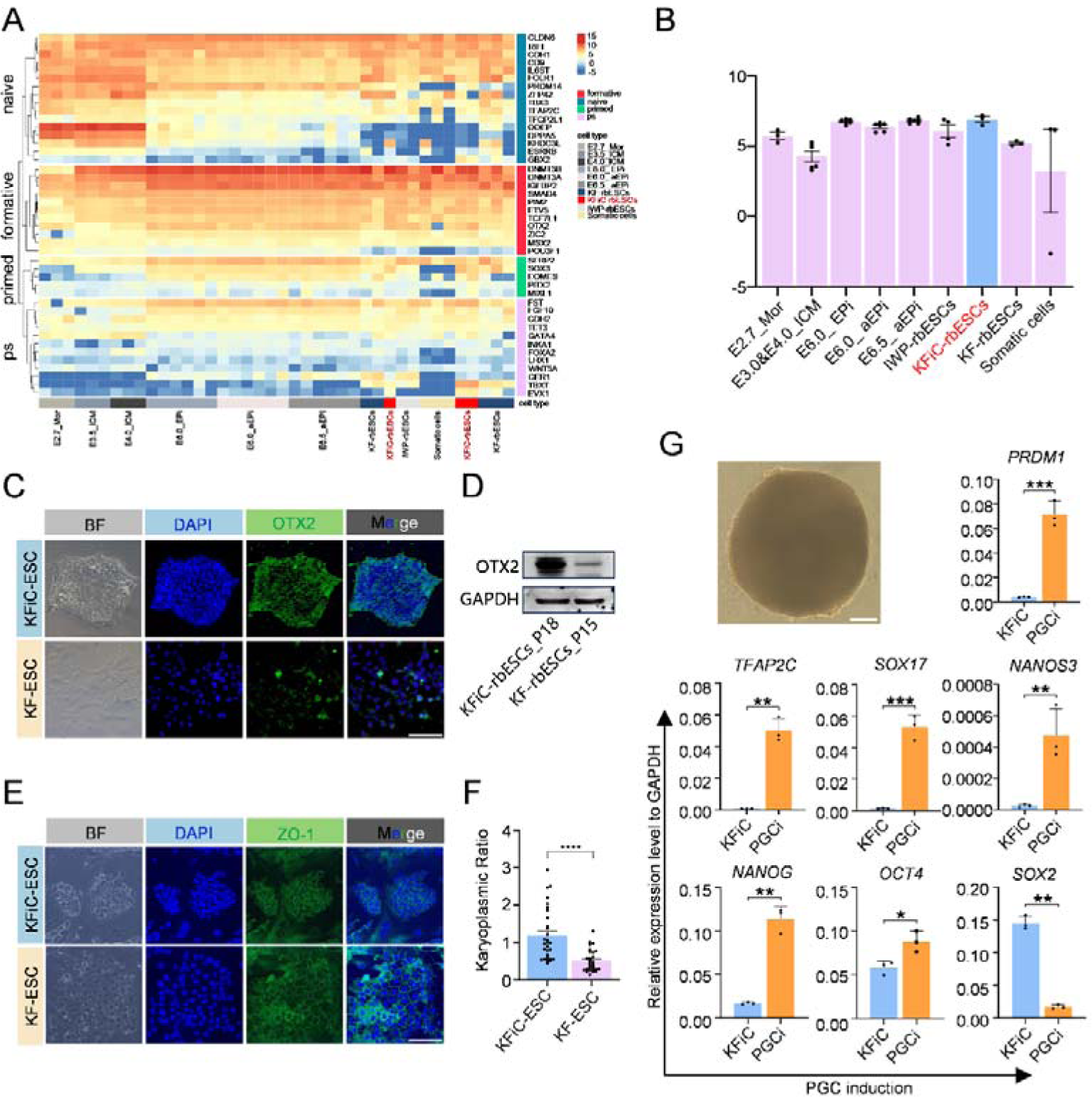
rbESCs Maintained in KFiC Express Formative Pluripotency Markers. (**A**) Hierarchical clustering of genes enriched in naïve, formative, primed, and primitive streak (PS) pluripotency states. Expression profiles are compared across embryonic stages (morula, E3.5 ICM, E4 ICM, E6 epiblast, E6 anterior epiblast, E6.5 anterior epiblast), somatic cells, and rbESCs cultured in KFiC, KF, 2i/LIF, or IWP. (**B**) OTX2 expression across selected samples. Expression levels are presented as log-transformed normalized counts per million (CPM). (**C**) Detection of OTX2 using immunofluorescence in P18 KFiC-rbESCs and P15 KF-rbESCs. Scale bar, 200 μm. (**D**) Validation of OTX2 expression at the protein level in KFiC-rbESCs and KF-rbESCs by western blot analysis. GAPDH was used as a loading control. (**E**) Immunofluorescence staining of ZO-1 in KFiC-rbESCs_P18 and KF-rbESCs_P15. Scale bar, 200 μm. (**F**) Nuclear-to-cytoplasmic (N/C) ratio analysis. Nuclear area was quantified from DAPI staining, and cytoplasmic area was delimited by ZO-1-positive cell boundaries; the nuclear-to-cytoplasmic (N/C) ratio was calculated from these areas. (**G**) Induction of KFiC-rbESCs toward rabbit primordial germ cell-like cells (rbPGCLCs). Schematic illustration of the rbPGCLC induction protocol, in which KFiC-rbESCs were cultured in GK15 induction medium supplemented with BMP4, SCF, EGF, and LIF. RT-qPCR analysis of pluripotency- and PGC-associated genes following differentiation of KFiC-rbESCs into rbPGCLCs. Data are presented as mean ±SD (n = 3). *P < 0.05, **P < 0.01, ***P < 0.001. Scale bar, 200 μm.

To assess the capacity of KFiC-rbESCs to differentiate toward the germline lineage, we followed a previously established protocol for PGCLC induction^38^. Following 6 days of differentiation, KFiC-rbESCs gave rise to three-dimensional PGCLC clusters and expressed key PGC-related markers, including *PRDM1*, *TFAP2C*, *SOX17*, and *NANOS3* (Fig. 3G). Concomitantly, the resulting PGCLCs retained *OCT4* and *NANOG* levels while displaying reduced *SOX2* expression (Fig. 3G), consistent with the transcriptional remodeling during PGC specification^34^.

Given distinct epigenetic and metabolic features of formative PSCs, we subsequently interrogated genes involved in these processes. DNMT3B, a de novo DNA methyltransferase involved in establishing DNA methylation patterns during naïve-to-formative pluripotency transition, was highly expressed from the ICM to the E6.5 epiblast and was also prominently expressed in KFiC-rbESCs (Fig. S3A). In addition, KFiC-rbESCs, together with in vivo epiblast populations, exhibited a glycolytic transcriptional signature characterized by elevated expression of *Pkm*, *Ldha*, and *Ldhb*^39^ (Fig. S3B). In contrast, genes related to mitochondrial oxidative phosphorylation (OXPHOS), including *NDUFV2*, *ATP5PD*, *UQCRQ*, *COX8A*, *CYTB*, and *ATP6*^35^, were comparatively reduced (Fig. S3B). Therefore, KFiC-rbESCs manifest a glycolytic metabolic signature resembling that of the pre-implantation epiblast rather than the OXPHOS-dominant metabolism observed in the ICM (Fig. S3B).

Together, these findings demonstrate that KFiC-rbESCs exhibit an integrated formative-like signature encompassing formative-associated gene expression, competence for PGCLC induction, epigenetic features, and glycolytic metabolism, supporting the establishment and maintenance of a stable formative-like state under KFiC conditions.

### Developmental competence of KFiC-rbESCs in rabbit embryos

We therefore evaluated the developmental potential of KFiC-rbESCs by introducing eGFP-labeled cells (KFiC-eGFP; Fig. S4A) into E1.5 rabbit morulae (Fig. 4A). At E5.0, KFiC-eGFP cells had survived and propagated extensively within chimeric blastocysts, with donor-derived cells broadly distributed throughout the blastocyst (Fig. 4B). More importantly, KFiC-eGFP cells were detected in both the ICM and trophectoderm (TE) compartments, with GFP signals observed within both lineages (Fig. S4B-I; Fig. S4C; Videos 1 and 2). At E6.0, KFiC-eGFP cells were also identified in embryonic outgrowths extruded from expanded chimeric blastocysts, further supporting their ability to persist and proliferate within the rabbit embryonic environment (Fig. S4B-II). We subsequently quantified donor-cell numbers in chimeric blastocysts and found that KFiC-eGFP cells increased more than twofold in most embryos during in vitro culture (Fig. 4C). Consistent with these observations, time-lapse imaging of chimeric embryos revealed sustained proliferation of KFiC-eGFP⁺ donor cells throughout embryonic development (Video 3).

**Figure 4.**
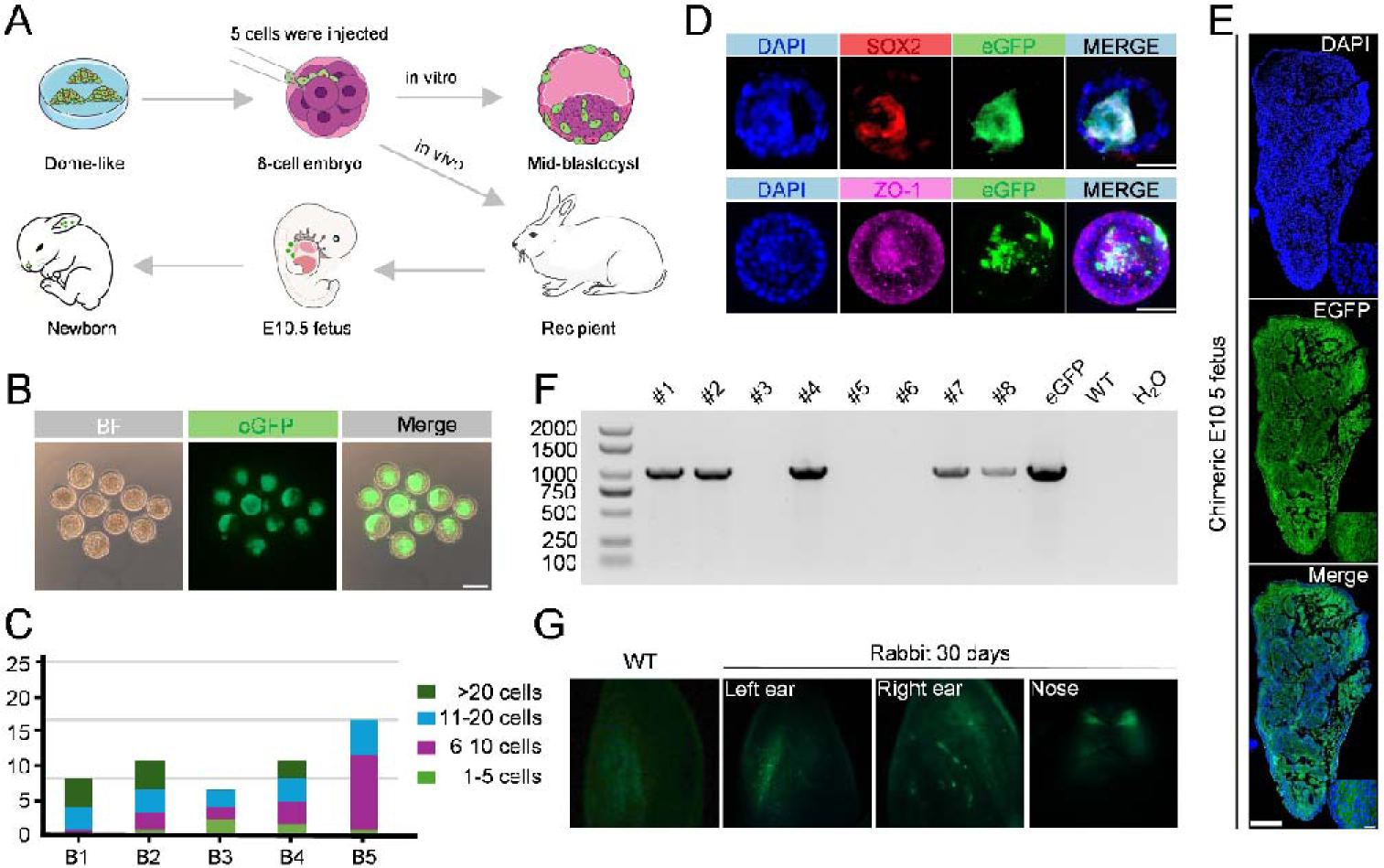
KFiC-rbESCs Contribute to Pre-implantation Embryos In Vitro and Embryogenesis In Vivo. (**A**) Schematic workflow of embryonic chimerism using KFiC-rbESCs. (**B**) Representative phase-contrast and GFP fluorescence images of chimeric rabbit embryos developed to the blastocyst stage in vitro. GFP signal indicates contribution of KFiC-rbESCs. Scale bar, 200 µm. (**C**) Quantification of GFP⁺ cells in chimeric embryos from five independent experiments. (**D**) Confocal microscopy of chimeric rabbit blastocysts (E4–E5.5) derived from early morulae (E2.8) microinjected with KFiC-eGFP cells. Scale bar, 200 µm. (**E**) Cross-sections of a morphologically abnormal E10.5 chimeric conceptus showing GFP-labeled KFiC-rbESCs (green) injected at the morula stage. Scale bar, 500 μm. (**F**) Gel electrophoresis of GFP transgene fragments amplified from genomic DNA extracted from the ears of eight full-term offspring. A GFP-containing plasmid served as the positive control, ear genomic DNA from a wild-type rabbit served as the negative control, and served as the no-template control. (**G**) Distribution of KFiC-rbESC-derived somatic cells (green) in one-month-old chimeric rabbit. Signals were detected in the ears and Nose. A rabbit derived from an embryo without KFiC-rbESC injection was used as a control.

To characterize the fate and developmental integration of KFiC-rbESCs in developing rabbit embryos, we examined their lineage specification and interaction with host cells in E4.0 chimeric blastocysts. Immunostaining for SOX2, an epiblast-specific protein, revealed that the majority of KFiC-eGFP cells were SOX2-positive, indicating that donor cells acquired or retained an epiblast-like identity within the host embryo (Fig. 4D). ZO-1 was detected at interfaces between neighboring KFiC-eGFP donor cells and host embryonic cells, suggesting the establishment of cell-cell junctions between donor and host cells (Fig. 4D).

We next transferred E2.8 chimeric embryos into surrogate females to evaluate the developmental capacity of KFiC-rbESCs during embryogenesis. Eight days later, one pregnant recipient was euthanized, and three E10.5 conceptuses were recovered. Among them, one morphologically degraded conceptus exhibited a high proportion of eGFP cells (Fig. 4E; Fig. S4D). Of the transferred embryos, eight developed to term, of which five were positive for the GFP transgene, as determined by PCR. Notably, one offspring displayed detectable eGFP⁺ donor-cell contribution in the ear and nasal tissue (Fig. 4F, G).

Together, these findings demonstrate that KFiC-rbESCs can efficiently integrate into developing rabbit embryos, proliferate extensively, and distribute broadly throughout the embryonic environment. The persistence of KFiC-rbESC-derived cells in a post-implantation conceptus and in tissues of full-term offspring further highlights their capacity to survive and differentiate during extended embryogenesis. Combined with their molecular features, these results support the characterization of KFiC-rbESCs as a formative-like pluripotent state with substantial early developmental competence.

### KFiC is able to establish ESCs with an intermediate pluripotent state in multiple mammalian animals

Given that rbESCs display formative-like pluripotency features in KFiC, we asked whether KFiC could similarly sustain ESCs from other mammalian species. We found that human ESCs cultured in 4CL (4CL-hESCs), mouse ESCs cultured in 2i/LIF (2i/LIF-mESCs), and pig ESCs cultured in EPSCM (EPS-pESCs) all survived and proliferated following transfer to KFiC. Strikingly, cells from all three species gradually adopted a flattened colony morphology similar to that of KFiC-rbESCs within 14 days of culture in KFiC (Fig. 5A). This morphological conversion was fully reversible, as KFiC-cultured cells regained their original colony morphology within 14 days after being returned to their respective culture conditions (Fig. 5A). These observations indicate that KFiC promotes a plastic intermediate pluripotent state across multiple mammalian species.

**Figure 5.**
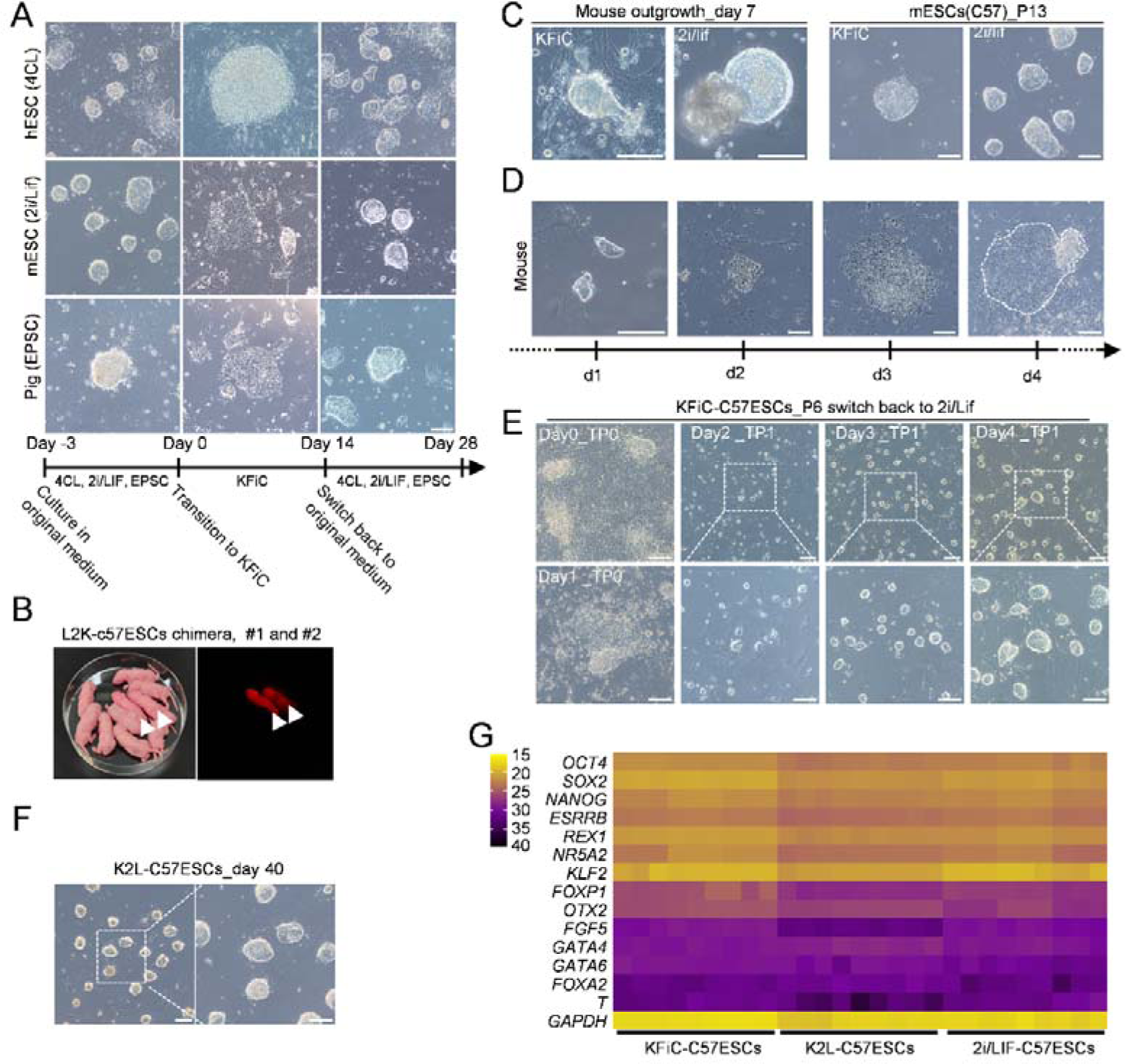
ESCs from Multiple Species Can Be Cultured in KFiC Medium. (**A**) Transient culture of established ESCs from human, mouse, and pig in KFiC medium for 14 days, followed by reversion to their respective original media for an additional 14 days. Original culture conditions: 4CL (Mazid et al.), 2i/LIF (Ying et al.), EPSCM (Gao et al.). Scale bar, 200 μm. (**B**) Red fluorescent chimeras (White arrow, offspring 1 and 2) were generated by injecting L2K-C57mESCs into ICR host blastocysts. (**C**) Morphology of C57 mouse blastocyst outgrowths and derived mESC lines established in KFiC or 2i/LIF medium. Scale bar, 200 μm. (**D**) Colony morphology of KFiC-C57mESCs over successive passages following single-cell dissociation with trypsin. Scale bar, 200 μm. (**E**) Phase-contrast images of C57mESCs undergoing naïve-like transition (K2L-mESCs). Cells were cultured in KFiC and then returned to 2i/LIF medium with 4 days for 1 passage. Images are shown at the indicated time points. Scale bar, 200 μm. (**F**) Morphology of naïve-transition C57-mESCs during serial passage in 2i/LIF medium. Scale bar, 200 μm. (**G**) RT-qPCR analysis of pluripotency state markers across cell lines. Expression of naïve, formative, and PS genes was measured in P8 KFiC-C57ESCs, P10 K2L-C57ESC (Swiched back to 2i/LIF in P6) and P8 2i/LIF-C57ESCs. Data from three independent experiments. Each dot represents a Ct value from one biological replicate.

To further evaluate the developmental potential of PSCs from other species maintained in KFiC, we injected L2K-C57ESCs (tdTomato-labeled 2i/LIF-derived C57 mouse ESCs cultured in KFiC for more than 24 days; Fig. S5A, 5B) into ICR mouse blastocysts and transferred the resulting embryos into recipient females. Among 18 full-term pups, three displayed extensive tdTomato fluorescence, indicating that mouse ESCs retained robust chimera-forming competence after prolonged culture in KFiC (Fig. 5B, Fig. S5C, D).

Given the stable proliferation and chimera-forming competence of mouse ESCs maintained in KFiC, we asked whether KFiC could support de novo establishment of mouse ESCs from blastocysts (KFiC-C57ESCs; Fig. 5C). Unlike 2i/LIF, in which each blastocyst predominantly gave rise to a single dome-shaped outgrowth, blastocysts cultured in KFiC produced two morphologically distinct types of outgrowths, characterized by a compact cluster emerging from the ICM and a surrounding monolayer of cells originating predominantly from the TE (Fig. 5C). Following establishment of a stable cell line from an ICM-derived outgrowth, KFiC-C57ESCs displayed morphological features similar to those of KFiC-rbESCs, rapidly forming compact dome-shaped colonies within 24 h after passaging and subsequently acquiring flattened EpiSC-like morphology over the following 72 h (Fig. 5D). RT-qPCR analysis further showed that KFiC-C57ESCs expressed high levels of formative-related markers *Otx2* and *Fgf5*, along with the naïve-associated markers *Esrrb*, *Rex1*, *Nr5a2*, and *Klf2*, consistent with an intermediate state between naive and primed pluripotency (Fig. 5G).

We further switched KFiC-C57ESCs to conventional 2i/LIF conditions, generating K2L-mESCs, which readily reacquired the dome-shaped colony morphology and molecular features of naïve mouse ESCs (Fig. 5E-G). Collectively, these findings demonstrate that KFiC can support a conserved intermediate pluripotent state across mammalian species and preserve the robust chimera-forming competence of mouse ESCs.

## Discussion

Our study establishes KFiC as a WNT/β-catenin signaling-modulated culture system capable of supporting the long-term maintenance of formative-like rabbit embryonic stem cells (rbESCs). KFiC-rbESCs retained competence for PGC induction and for contribution to embryonic development following introduction into rabbit morulae, highlighting their developmental plasticity and providing a foundation for developing ESC-based approaches to generate genetically modified rabbit models. Intriguingly, KFiC was also capable of stabilizing a reversible intermediate pluripotent state in PSCs from multiple mammalian species, including mouse, human, and pig. In mice, KFiC-adapted ESCs retained high developmental competence, as evidenced by their robust contribution to chimeric embryos, further supporting the ability of KFiC to modulate pluripotency toward an intermediate state along the naïve-to-formative continuum. These findings provide motivation for further optimization of KFiC toward establishing more naïve-like and developmentally competent rbESCs.

Previous studies have established FTW as a cross-species culture system capable of stabilizing formative-like pluripotent stem cells^28^, which exhibit several hallmarks, including upregulation of *Otx2*, epigenetic remodeling characterized by increased H3K4me3 and reduced H3K27me3, enhanced glycolytic metabolism, and competence for PGCLC induction and chimera formation. Although FTW supported the formation of dome-shaped outgrowths from rabbit blastocysts, an effect that was further enhanced by supplementation with 5% KOSR, rbESCs cultured in FTW with or without 5% KOSR underwent progressive differentiation and ultimately failed to sustain long-term self-renewal. Unlike the canonical concentration of CHIR99021 used in mouse ESC culture, Chen et al.^30^ and Li et al.^31^ reported that reducing the CHIR99021 concentration from 3 μM to 1 μM in 2i/LIF medium in rat ESCs (T2iL) attenuates GSK3 inhibition, thereby decreasing nuclear β-catenin accumulation and preventing excessive activation of the WNT/β-catenin pathway. This modulation suppresses the expression of downstream differentiation-associated genes such as CDX2, enabling sustained self-renewal of rat ESCs and facilitating their application in tetraploid complementation assays. In our hands, however, both FTW and T2iL were insufficient to support the long-term culture of rbESCs, suggesting that simple activation of WNT/β-catenin signaling through CHIR99021 is not sufficient to sustain rbESC pluripotency. In contrast, KFiC coordinately modulates WNT/β-catenin signaling through simultaneous activation by CHIR99021 and inhibition by IWR-1, thereby constraining excessive nuclear accumulation of β-catenin and promoting its predominantly cytoplasmic localization^14^, suggesting that a tuned level and subcellular distribution of WNT/β-catenin signaling may be critical for efficient derivation and long-term maintenance of rbESCs.

More recently, Yang et al. reported a culture system termed FACE that supports the derivation and long-term maintenance of ESCs with formative-like characteristics from ruminant species ^29^. In our results, however, although FACE supported the derivation of dome-shaped outgrowths, it was insufficient to sustain long-term self-renewal of the derived cells, similar to FTW. Notably, FACE and KFiC share a comparable strategy for WNT modulation by combining the WNT activator CHIR99021 and the inhibitor IWR-1, yet their capacities to sustain rbESCs differed markedly. A key difference between the two media is that KFiC contains FBS in its basal medium, whereas FACE does not. This difference may contribute to the superior capacity of KFiC to support the long-term maintenance of rbESCs. Consistent with this possibility, replacing FBS with KOSR in KFiC resulted in differentiation of all tested rbESC lines despite preservation of the CHIR99021/IWR-1 combination (data not shown). This observation is in line with previous rbESC culture systems, including KF and bXC, in which serum-containing conditions supported sustained proliferation and maintenance of characteristic colony morphology^4,8^, further suggesting that serum-derived factors may play pivotal roles in sustaining rbESC self-renewal.

Furthermore, a recent E4/6i culture system was developed by incorporating six pathway inhibitors to enable the derivation and maintenance of ESCs across multiple species, including the rabbit^9^. Nevertheless, rbESCs acquired chimera-forming competence only after the addition of the LATS inhibitor TDI-011536 (TDI), resulting in the E4/7i condition, and even then, chimeric contribution remained relatively inefficient. These findings suggest that robust developmental competence of rbESCs may be governed by regulators whose requirements differ from those of ESCs from other mammalian species.

In our study, KFiC also supported PSCs from multiple species. Notably, mouse ESCs established or adapted to KFiC readily reacquired a naïve pluripotent phenotype upon transition to 2i/LIF, indicating that KFiC may maintain a metastable intermediate state positioned between naïve and primed pluripotency^40,41^. The reversibility of this transition suggests that KFiC does not irreversibly restrict cells toward a primed state but instead preserves pluripotent plasticity that can be redirected toward naïve pluripotency under appropriate conditions. Whether KFiC-rbESCs similarly occupy an intermediate pluripotent state, as observed for KFiC-C57ESCs, remains to be determined. Still, this possibility provides a useful conceptual and cellular framework for future efforts to further refine KFiC conditions toward establishing authentic naïve rbESCs.

Importantly, although KFiC-rbESCs exhibited extensive developmental competence during the early stages of embryonic development, their contribution to post-implantation and full-term development remained limited. A recent study demonstrated that forced resetting toward a naïve pluripotent state by introducing naïve-associated factors can endow rabbit iPSCs with germline chimera-forming competence^10^. However, such factor-based reprogramming approaches may have limitations for generating disease models because sustained or ectopic expression of factors such as Eras has been associated with tumorigenic potential^42^. In this study, we recovered one morphologically abnormal E10.5 conceptus with a high proportion of KFiC-rbESC-derived cells, suggesting that KFiC-rbESCs may possess broad developmental potential and can contribute extensively to embryonic tissues, but their developmental state may not yet be fully compatible with sustained post-implantation development. One possible explanation is a developmental asynchrony between the donor KFiC-rbESCs and the recipient embryo. Synchronization between the developmental state of donor PSCs and that of the host embryo is critical for donor-cell proliferation, lineage integration, and subsequent differentiation^43^. Unlike mouse ESCs, which are maintained in a naïve pluripotent state and readily integrate into embryos at various pre-implantation stages, including 2-cell stage^44^, morulae^45^, and blastocysts^46^, KFiC-rbESCs exhibit formative-like characteristics and may therefore correspond to a later epiblast stage that is not optimally matched to the recipient morula used in this study. These species-specific differences in PSC-embryo interactions may contribute to developmental asynchrony through differences in signaling responsiveness, cell-cycle state, and lineage restriction, thereby limiting the ability of formative-stage rbESCs to integrate efficiently into the earlier embryonic environment. In this regard, introducing KFiC-rbESCs into developmentally matched later-stage embryos, such as expanded blastocysts recovered in utero, might provide an alternative strategy to enhance donor-cell contribution and ultimately facilitate the generation of germline-contributing chimeric rabbits in future studies.

## Materials and Methods

### 1. Animals

New Zealand White rabbits were purchased from the Guangdong Medical Laboratory Animal Center (Foshan, China) and maintained at the rabbit facility (clean grade) of Guangzhou Institutes of Biomedicine and Health (GIBH), Chinese Academy of Sciences. During the animal study, the rabbits were given sufficient food and water at 25 ±1 °C under a 12 h light/dark cycles. Over 5-month-old male and female rabbits were used in subsequent experiments. Mouse was purchased from BesTest (Zhuhai, China). They were randomly divided into the individually ventilated cages and were given sufficient food and water at 25 ±1 °C. Over 8-weeks old male and female mice were used in follow-up experiments. All experiments and their operations were approved by the Institutional Animal Care and Use Committee of GIBH.

### 2. Production and collection of rabbit and mouse Embryos

Sexually mature female rabbits were superovulated by subcutaneous administration of follicle-stimulating hormone (FSH), pregnant mare serum gonadotropin (PMSG) and human chorionic gonadotropin (hCG), then mated with fertile or castrated males. At 22 hours post-coitum, zygotes were flushed from the oviductal fimbriae using PBS supplemented with 5% FBS. Recovered embryos were collected with a mouth-operated pipette and washed three times in EBSS, placed under paraffin oil, and cultured in EBSS until they reach the 8-cell or blastocyst (E4) stage.

Mouse embryos were obtained from 8-week-old females treated with PMSG and hCG following natural mating. Zygotes were collected from oviducts 16 h post-hCG in M2 medium, then cultured in KSOM droplets at 37 °C with 5% CO₂ to the blastocyst stage.

### 3. Derivation and culture of rabbit and mouse pluripotent stem cell lines

Undifferentiated rabbit embryonic stem cells (rbESCs) were established and maintained on mitomycin C-inactivated mouse embryonic fibroblasts (MEFs) in KFiC culture medium, which consisted of DMEM/F12 supplemented with 10% fetal bovine serum (FBS), 10% KnockOut Serum Replacement (KOSR), 1× GlutaMAX, 1× non-essential amino acids (NEAA), 50 μM 2-mercaptoethanol, 1% penicillin-streptomycin (10,000 U/mL each), 8 ng/mL recombinant human FGF2 (rhFGF, Peprotech, 100-18B-100) (named KF basic medium), 1 μM CHIR99021 (Selleck, S1263), and 1 μM IWR-1 (Selleck, S7086).

For rbESC and KFiC-mESC line derivation, E4 blastocyst-stage rabbit embryos were directly seeded into 24-well plates containing feeder layers and cultured in various medium including KFiC, KFF, 2i/LIF, FTW and 4CL at 38.5 °C under 5% CO₂. After approximately 6 days, blastocyst outgrowths were dissociated with 0.05% trypsin and passaged onto fresh feeder cells in 24-well plates. The culture medium was refreshed daily and visible embryonic stem cell colonies typically appeared within 3–4 days and were subsequently serially passaged under the same conditions to establish stable rbESC lines.

E3.5 mouse blastocysts were seeded onto feeder layers in 24-well plates and cultured in KFiC medium at 37 °C with 5% CO₂. After 6 days, outgrowths were dissociated with 0.05% trypsin and passaged onto fresh feeders. Medium was refreshed daily. Visible ES cell colonies emerged within 3–4 days and were serially passaged to establish stable KFiC-mESC lines. Regular testing confirmed the absence of mycoplasma contamination.

### 4. TOPFlash assay

The TOPFlash reporter plasmid (Beyotime, A0181) was electroporated into KFiC-rbESCs and KF-rbESCs. Transfected cells were seeded onto feeder layers in 6-well plates. After 72 h, luciferase activity was measured using the detection kit (Beyotime, RG005) and relative luminescence units (RLU) were quantified.

### 5. Vector construction and acquisition of the eGFP⁺ and RFP⁺ cell lines

eGFP⁺ and RFP⁺ cells were generated by electroporating KFiC-rbESCs and KFiC-mESCs-C57 with PiggyBac (PB) transposon vectors encoding pCAGGS-EGFP-T2A-PURO or pCAGGS-tdTomato-T2A-PURO, together with a PB transposase expression plasmid. Following the emergence of single-cell-derived colonies, stable eGFP⁺ or RFP⁺ colonies were selected using 1 μg/mL puromycin. Clones exhibiting the highest eGFP or RFP fluorescence were subsequently isolated by manual picking with glass capillaries and expanded to establish rabbit KFiC-eGFP⁺ and mouse KFiC-RFP⁺cell lines. The genotyping primers for eGFP⁺ cells are eGFP-F: gagccgcagccattgccttt and eGFP-R: gggtctttgctcagggcgga. Cells at passages 15-20 were used for microinjection; both lines were maintained for >60 passages without loss of fluorescence or pluripotency.

### 6. Cell microinjection, embryo culture and transfer

In vivo-produced embryos were placed in EBSS droplets under paraffin oil and cultured at 38.5 °C with 5% CO₂ until the 8-cell stage (E1.5). At this stage, each embryo was injected with 5-10 eGFP⁺ cells using a micromanipulation system. Then, the embryos were placed in droplets of a 1:1 KFiC:EBSS mixture for further cultivation. When the chimeric embryos developed to E2.8, the embryos were transferred into the oviducts of pseudopregnant females through the fimbriae using a transfer pipette. On the 8th day after transplantation, the implanted embryos (E10.5) were obtained from the uterine horn, or the full-term offspring were obtained on days 29-30 after transplantation.

For the mouse chimerism experiment, ICR mouse embryos that had developed into blastocysts in the M2 were injected with 10-15 RFP⁺ KFiC-mESCs, and then the embryos were transplanted into the uteri of pseudopregnant recipients. Seventeen days later, pups were obtained.

### 7. Immunofluorescence analysis of cells and embryos

rbESCs were cultured in 4-well plates. When distinct colonies emerged, the cells were washed three times with PBS and fixed in 4% PFA at room temperature for 30 min. E4.0 or E4.5 chimeric embryos were similarly fixed in 4% PFA for 30 minutes. After fixation, both rbESC colonies and embryos were washed three times with PBS (3 min per wash). They were then permeabilized and blocked in PBS containing 0.3% Triton X-100 and 10% goat serum for 1 hour at room temperature. Subsequently, samples were incubated overnight at 4°C with primary antibodies against SOX2 (Cell Signaling Technology, #4900S, 1:200), OCT4 (BD Biosciences, 611202, 1:20), OTX2 (R&D Systems, AF1979-SP, 1:50), and GFP (Invitrogen, A10262, 1:500). The next day, samples were washed four times with PBST (PBS containing 0.05% Tween-20), 3 min each wash, and then incubated with appropriate secondary antibodies—488 goat anti-mouse (Proteintech, RGAM002, 1:1000), 555 goat anti-mouse (Proteintech, RGAM003, 1:1000), and 488 goat anti-chicken (Abcam, ab150173, 1:1000)—for 1 hour at room temperature. Then, samples were again washed four times with PBST (3 minutes per wash). Nuclei were counterstained with DAPI (Beyotime, C1002, 1:1000) for 8 minutes. Finally, rbESC colonies were mounted in 30% glycerol, and embryos were preserved in 30% glycerol droplets for imaging. Without prior fixation, fresh E10.5 embryos were embedded in OCT compound and cryosectioned at a thickness of 25 μm. The sections were then used directly for staining.

### 8. Embryoid body (EB) formation and teratoma formation

Undifferentiated KFiC-rbESCs were dissociated with 0.05% trypsin and subjected to feeder depletion by incubating the cell suspension in a 10 cm cell culture dish coated with 0.2% gelatin for 15 minutes in KF medium. The non-adherent cells in the supernatant were collected by centrifugation, resuspended in KF medium, and then seeded at 1 × 10⁴ cells per well in ultra-low attachment U-bottom 96-well plates. After 7 days of culture, embryoid bodies (EBs) formed.

For teratoma formation, undifferentiated KFiC-rbESCs were dissociated with 0.05% trypsin and resuspended in a 1:1 mixture of DMEM/F12 and PBS. A total of 1 × 10⁷ cells in 150 μl of suspension were injected subcutaneously into the bilateral inguinal regions of 8-week-old nude mice. Approximately 7 weeks post-injection, teratomas had reached a size of about 2 cm³. The mice were euthanized and teratomas were harvested for H&E staining and bulk RNA sequencing.

### 9. Quantitative reverse transcription PCR

Total RNA was extracted from cells using Trizol (Servicebio, G3013) reagent. For cDNA synthesis, 2 μg of total RNA was reverse-transcribed using the PrimeScript™ RT Reagent Kit with gDNA Eraser (TaKaRa, RR047A), and the resulting cDNA was diluted 10-fold as PCR template or stored in −80℃. Quantitative real-time RT-PCR (RT-qPCR) was performed on a CFX96 system (Bio-Rad) using 1 μL of diluted cDNA and SYBR® Green Premix Ex Taq™ (Vazyme, Q711) according to the manufacturer’s instructions. Gene expression levels were normalized to GAPDH as an internal control. The primer sequences used are listed in Table S1.

### 10. Western Blot Analysis

KFiC-rbESCs and KF-rbESCs were lysed in radio immunoprecipitation assay (RIPA) buffer (Beyotime, Shanghai, China). After centrifugation, the supernatant containing total protein was collected, and the protein concentration was determined by NanoDrop. The protein was separated by 12% sodium dodecyl sulfate-polyacrylamide gel electrophoresis (SDS-PAGE), and then transferred to PVDF (Millipore, Darmstadt, Germany) membrane. After blocking with 5% skim milk for 2 h, membranes were incubated overnight at 4 ℃ with primary antibodies against SOX2 (1:1000; Cell Signaling, 4900s), OCT4 (1:1000; BD Biosciences, 611202), OTX2 (1:1000; R&D systems, AF1979-SP), and GAPDH (1:1000; HUABIO, ER1901-65). Then, the PVDF membrane was washed with Tris-buffered saline with Tween-20 (TBST) solution for 4 times, each time for 5 min, and then the PVDF membrane was incubated with secondary antibody goat anti-rabbit antibody (1:1000), goat anti-mouse antibody (1:1000) (Santa Cruz, California, USA) or donkey anti-goat (1:1000) (Beyotime, A0181) at room temperature for 1 hour, then the PVDF membrane was washed with TBST solution for 4 times, each time 5 min. According to the manufacturer’s instructions, the specific protein bands were detected with the Enhanced Chemiluminescence (ECL) Detection Kit (Beyotime).

### 11. PGC-like cell (PGCLC) induction of KFiC-rbESCs

When KFiC-rbESCs reached approximately 70–80% confluency, cells were dissociated into single-cell suspensions using 0.05% trypsin. The dissociated cells were seeded onto 0.2% gelatin-coated dishes for feeder cell removal. Subsequently, approximately 5,000 cells were seeded per well into 96-well U-bottom ultra-low attachment plates (Corning, 7007) in 100 μl of PGCLC induction medium and cultured for 6 days until characteristic PGCLCs morphology was observed. The PGCLC induction medium consisted of GK15 basal medium supplemented with 200 ng/mL BMP4 (R&D Systems, 314-BP-010), 100 ng/mL SCF (R&D Systems, 455-MC-020), 50 ng/mL EGF (R&D Systems, 236-EG-200), 10 ng/mL human LIF (Merck Millipore, LIF1005), and 10 μM ROCK inhibitor (MCE, HY-10583). GK15 medium was composed of GMEM (Gibco, 11710-035) supplemented with 15% KOSR, 2 mM L-GlutaMAX, 1× NEAA, 1 mM sodium pyruvate, 100 μM 2-mercaptoethanol, and 1% penicillin–streptomycin solution (10,000 U/mL penicillin and 10,000 U/mL streptomycin). On day 3 of induction, 50 μl of PGCLC induction medium was added to each well. PGCLC-like cells were collected on day 6 for subsequent analyses.

### 12. RNA-sequencing and data processing

Total RNA was extracted using TRIzol™ Reagent (Thermo Fisher, 15596018), and strand-specific RNA-seq libraries were prepared with the NEBNext® Ultra™ II Directional RNA Library Prep Kit for Illumina (NEB, E7760) according to the manufacturer’s instructions. All raw RNA-Seq reads were filtered using fastp^47^ with the following parameters: fastp -w 16 -z 4 -q 20 -u 30 -n 10. Filtered reads were aligned to the rabbit reference genome (mOryCun1.1, GCF_964237555.1) using HISAT2^48^ with default parameters. Potential duplicate alignments were removed using sambamba^49^ with: sambamba markdup -r -t 24 −l 5. Cleaned BAM files were further analyzed using the R packages Rsubread^50^ and edgeR. Gene-level read counts were obtained with feature Counts using the following parameters: feature Counts (GTF.featureType = “exon”, GTF.attrType = “gene_id”, isPairedEnd = TRUE, strandSpecific = 2). Normalization and differential expression analysis were performed with edgeR, where genes with low expression were filtered prior to analysis and normalization was carried out via the median-of-ratios method. Hierarchical clustering heatmaps for global transcriptomes and lineage-specific genes were generated using the ComplexHeatmap package^51^. Scatter plots were produced with ggplot2^52^, and pathway and Gene Ontology (GO) enrichment analyses were performed using DAVID^53,54^. In addition, raw sequencing data from public repositories with accession numbers E-MTAB-10892^34^, E-MTAB-10893^34^, PRJNA529333^55^, and PRJNA743177^35^ were downloaded and processed through the identical pipeline described above.

### 13. Quantification and statistical analysis

All data are presented as the mean ± SD, with detailed statistical parameters provided in the corresponding figure legends. GraphPad Prism 8 (GraphPad Software, USA) was used for data visualization and statistical analysis. For comparisons among multiple groups, one-way ANOVA followed by appropriate post hoc tests was employed. Statistical significance was defined as P < 0.05. Significance levels are indicated as follows: *P < 0.05, **P < 0.01, ***P < 0.001 and ****P < 0.0001.

