## Supplementary Figure1-5 for "Establishment of Rabbit Formative-like Pluripotent Cells by Bidirectional Modulation of WNT Signaling Pathway"

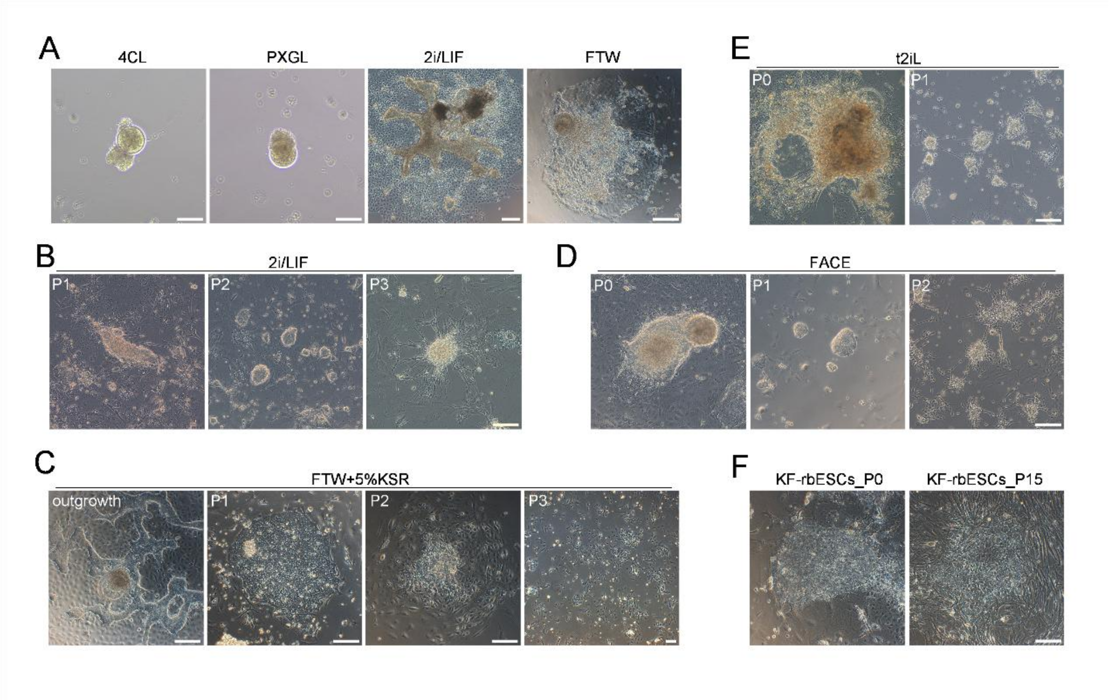


**Supplementary figure 1. Optimization of culture regimens for rbESCs.**

A. Rabbit blastocysts cultured under previously reported conditions for 6 days. 4CL (rhLIF, PD0325901, DZNep, TSA, IWR-1, Activin A, L-ascorbic acid), PXGL (PD0325901, XAV939, Gö6983, hLIF), 2i/LIF (CHIR99021, PD0325901, mLIF), FTW (bFGF, Activin A, CHIR99021). Scale bar, 200μm.

B. Morphology of rabbit blastocyst derivatives cultured in 2i/LIF medium. Scale bar, 200μm.

C. Morphology of outgrowths and their derivatives cultured in FTW medium supplemented with 5% KOSR. Scale bar, 200μm.

D. Morphology of rabbit blastocyst derivatives cultured in FACE medium. FACE (bFGF, Activin-A, IWR-1, CHIR99021). Scale bar, 200μm.

E. Morphology of rabbit blastocyst derivatives cultured in T2iL (CHIR99021, PD0325901, mLIF ) medium. Scale bar, 200μm.

F. rbESCs cultured in KF medium, as described by Osteil et al. Scale bar, 200μm.

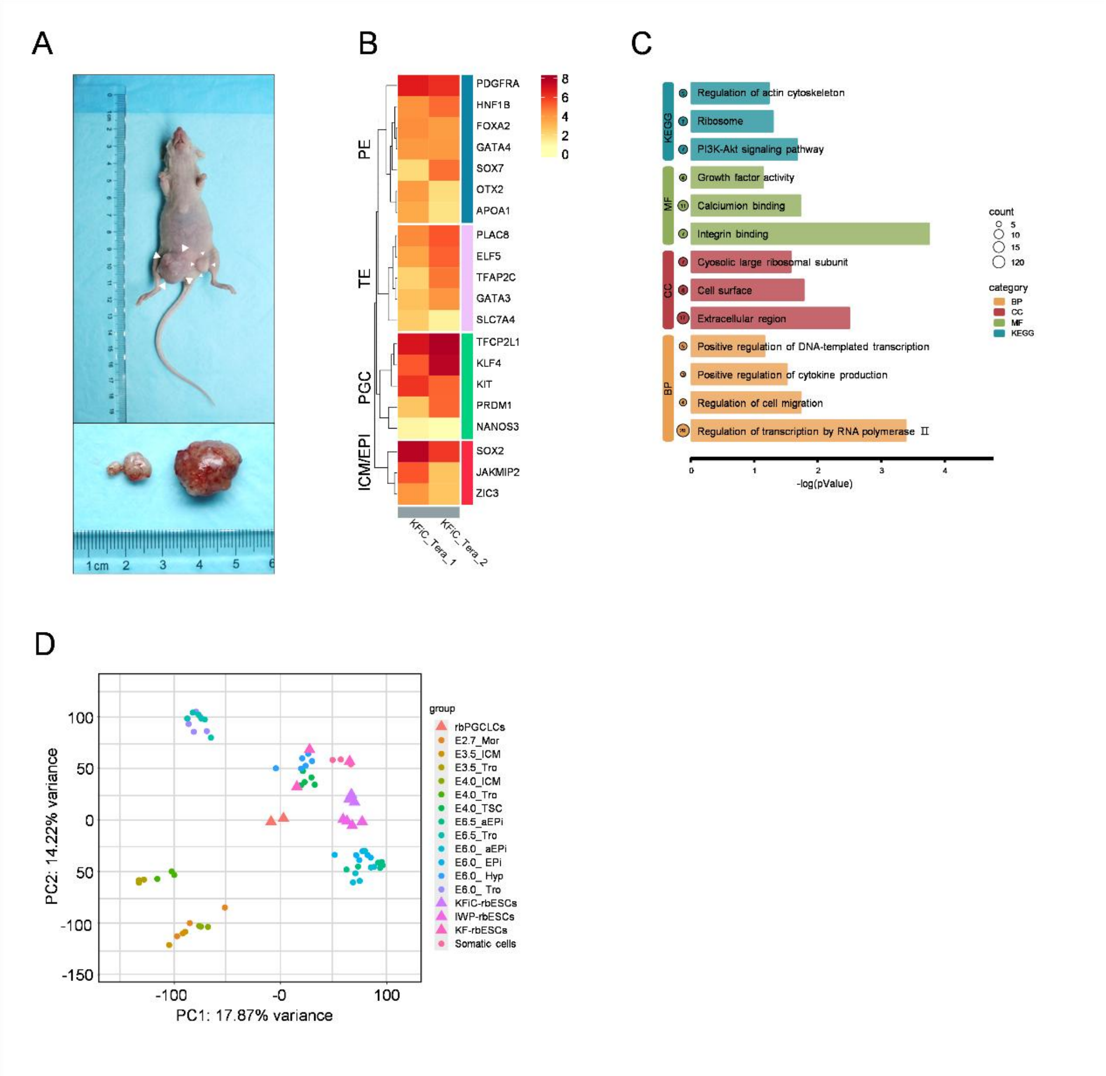


**Supplementary figure 2. Pluripotency and Transcriptomic Profiles of rbESCs Cultured in KFiC Medium.**

A. Teratoma produced by injection of P18 KFiC-rbESCs into immunodeficient mice.
B. Heatmap of germ-layer marker gene expression in two independently derived teratomas, assessed by bulk RNA-seq .
C. Gene Ontology (GO) enrichment analysis of the top 200 differentially expressed genes (DEGs) between KFiC-rbESCs and IWP-rbESCs. Significantly enriched terms were defined using a Benjamini–Hochberg adjusted p-value < 0.05.
D. Projection of KFiC-rbESCs and KF-rbESCs onto rabbit embryonic single-cell transcriptomic data using PCA. Rabbit single-cell data were obtained from W. Bouchereau et al., 2022.

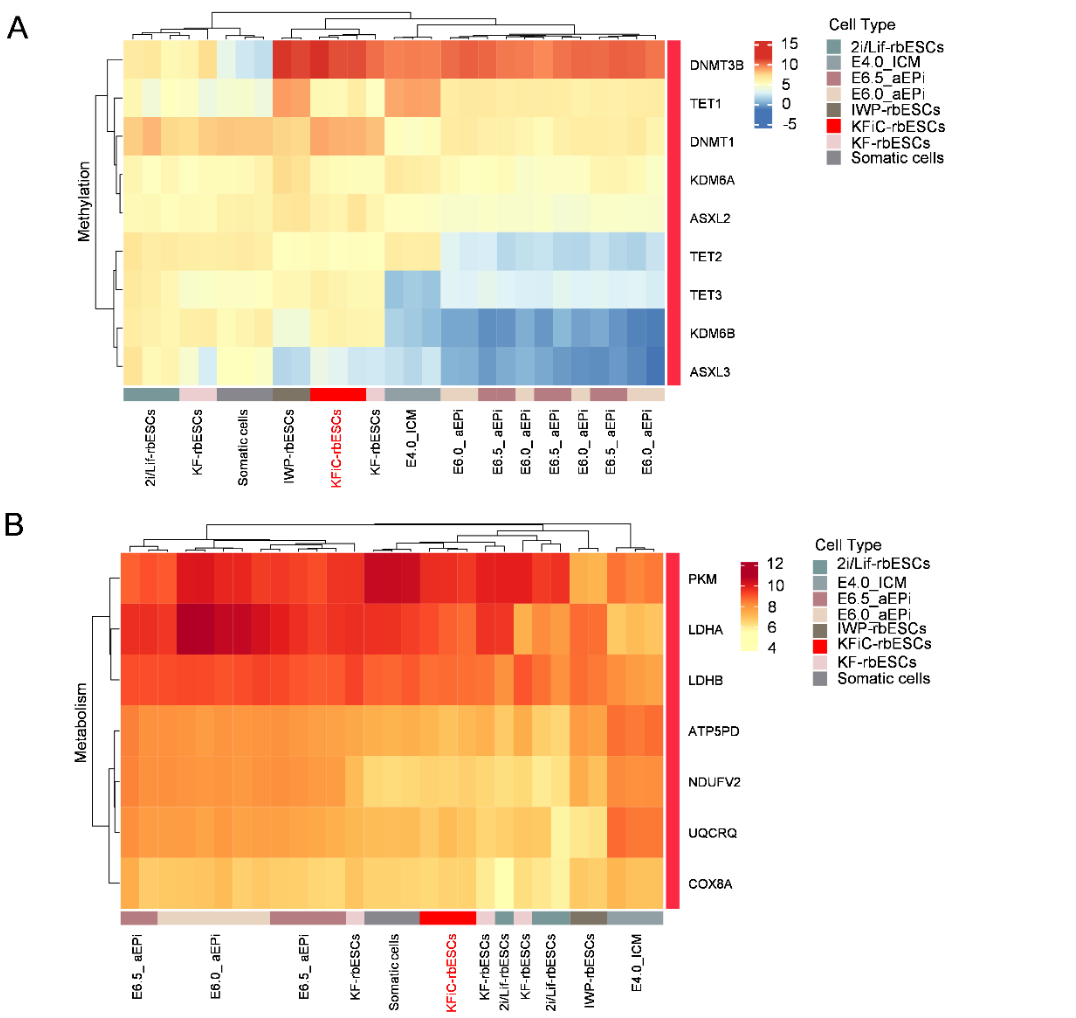


**Supplementary figure 3. Transcriptomic profiling defines KFiC-rbESCs as a formative-like epiblast model.**

A. Heatmap of genes encoding DNA methylation regulators across morula, E4_ICM, E6_aEPi, E6.5_aEPi, somatic cells, KF-rbESCs, 2i/LIF-rbESCs, IWP-rbESCs, and KFiC-rbESCs.
B. Heatmap clustering of metabolism enriched genes between morula, E4_ICM, E6_aEPi, E6.5_aEPi, somatic cells, KF-rbESCs, 2i/LIF-rbESCs, IWP-rbESCs and KFiC-rbESCs terms.

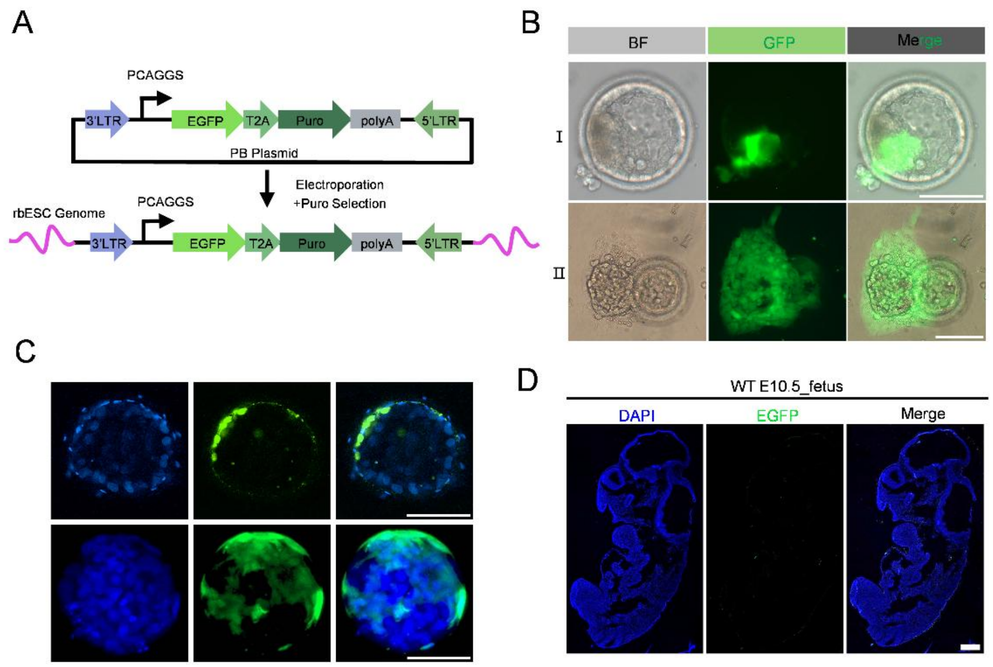


**Supplementary figure 4. Extensive In Vitro Proliferation of KFiC-rbESCs in Chimeric Rabbit Embryos.**

A. Schematic of eGFP-Containing PB Transposon Vector Electroporation into KFiC-rbESCs.

B. Representative phase-contrast and GFP fluorescence images of chimeric rabbit blastocysts developed in vitro to the blastocyst stage. (I) Enlarged view showing colonization of the ICM by KFiC-eGFP$^{+}$ cells. (II) In late blastocysts, KFiC-eGFP$^{+}$ cells extensively proliferated, emerged from the embryo proper, and contributed to outgrowths, recapitulating early post-implantation morphogenesis in vitro . Scale bar, 200 µm.

C. Confocal images of chimeric rabbit blastocysts developed in vitro to the blastocyst stage (E4-E4.5) following microinjection at 8-cell stage (E1.5) with KFiC-eGFP$^{+}$ cells. Scale bar, 200 µm.

D. Confocal images of E10.5 WT rabbit conceptus. Scale bar, 500 µm.

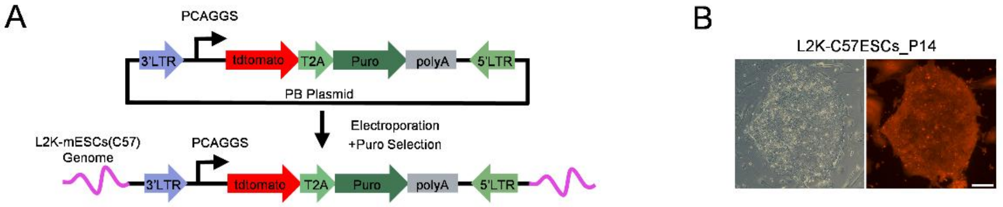


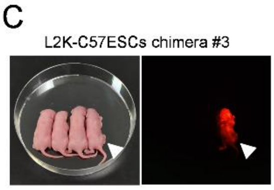


D

| Cell line | No. of embryos transferrd (ET) | No.of pups at full-term(% of ET) | No. of chimeras | Chimeras frequency (% of full-term) |
|---|---|---|---|---|
| L2K-C57ESCs | 37 | 14(37.8) | 3 | 21.4 |

**Supplementary figure 5. ESCs from Multiple Species Can Be Cultured in KFiC Medium.**

A. Schematic of tdtomato-Containing PB Transposon Vector Electroporation into L2K-C57ESCs.

B. Representative images showing a colony of P14 L2K-C57ESCs. Scale bar, 200μm.

C. Red fluorescent chimeras (White arrows, offspring 3) were generated by injecting L2K-C57ESCs into ICR host blastocysts.

D. Summary of blastocyst injections using L2K-C57ESCs.
